# XIAP-associated factor 1 protects against viral-induced neuropathogenesis

**DOI:** 10.64898/2026.05.26.727868

**Authors:** Rodrigo Cañas-Arranz, Melissa Uccellini, Fahmida Alam, Soner Yildiz, Rocío Seoane, Sara S. El Zahed, Adolfo García-Sastre

## Abstract

XIAP-associated factor 1 (XAF1) is a proapoptotic protein known to be involved in tumor suppression and regression whose gene expression has been reported to be dysregulated in a wide variety of tumor malignancies by different molecular mechanisms. Using a sterile alpha and TIR motif containing 1 (SARM1) knockout mouse model, we previously showed that XAF1 could be a candidate gene for protection against neurotropic virus infection. Here, using a CRISPR-knockout XAF1 mouse model, we show that XAF1 knockout mice are more susceptible to disease after VSV infection, a well-known neurotropic virus in mice. Interestingly, VSV-increased sensitivity in XAF1 knockout mice was not accompanied by differences in viral replication in the central nervous system (CNS). Nevertheless, infection of XAF1 knockout mice resulted in an increased pro-inflammatory response and immune cell infiltration into the CNS compared to that in wild-type mice. Similarly, XAF1 knockout mice showed slight increase to disease after infection with a different neurotropic virus, West Nile Virus (WNV). However, no differences in viral disease due to the absence of XAF1 were found upon infection with a respiratory virus such as influenza A virus (IAV). *In vitro*, XAF1-deficient cells showed a significant increase in interferon-stimulated genes (ISGs) expression upon stimulation with IFN and with different PAMPs, such as Poly(I:C), HT-DNA and LPS. Consistently, ectopic overexpression of XAF1 decreased IFN-signaling in a dose-dependent manner. Altogether, the data presented here suggest that the host factor XAF1 has a protective role in viral-induced neuropathogenesis due to excessive IFN responses.

**Author summary:** We previously identified XIAP-associated factor 1 (XAF1) as a candidate cell factor involved in viral phenotypes attributed to SARM1 deficiency. Even though the role of this factor has been studied in the cancer field as a proapoptotic tumor suppressor, its relevance in the context of viral infections has remained unclear. Here, we show that XAF1-deficient mice show increased susceptibility upon neurotropic virus infection and augmented levels of proinflammatory cytokines. We observe higher immune cell infiltration into the brain and disease exacerbation upon infection in mice lacking XAF1. This increased pathology is restricted to the brain, since no morbidity was observed upon infection with a respiratory virus, such as influenza virus. Gene expression analysis unveiled an unbalanced immune response in XAF1-deficient mice resulting in an elevated proinflammatory response and diminished capacity to restore homeostasis. Our data demonstrates the protective capacity of XAF1 and provides new insights into the host response against virus infections.

## Introduction

X-linked inhibitor of apoptosis (XIAP)-associated factor 1 (XAF1) is a proapoptotic protein involved in tumor regression whose gene expression is increased by IFN (1). XAF1 domains include a TNF receptor-associated factor (TRAF)-like N-terminal domain, a middle domain and a XIAP-interacting domain at the C-terminal domain (2). Initially identified from cancer studies, it has been observed that tumor cells have dysregulated XAF1 levels due to hypermethylation of its promoter (3). Short-truncated isoforms of XAF1 derived from alternative splicing of XAF1 RNA are upregulated in transformed cells, and it has been suggested that these shorter isoforms play a dominant negative function. The molecular mechanisms by which XAF1 induces apoptosis have been previously studied (1, 4-6).

XAF1 was initially described as an apoptotic protein promoting cell death by binding and inhibiting XIAP (7). However, pro-apoptotic activity of XAF1 has been described in XIAP-deficient cells, suggesting additional mechanisms by which can promote apoptosis. For instance, XAF1 is a transcriptional target of p53 and displaces MDM2 from p53, promotes p53 phosphorylation and degrades p21, leading to apoptosis (6). Also, a role of XAF1 has also been described in a XIAP-independent mechanism to interfere with the AKT signaling pathway, suggesting additional roles of this factor in cell metabolism and proliferation (8).

Since IFN positively regulates XAF1 expression, it has been suggested that XAF1 could play a role in IFN-mediated antiviral immunity and inflammation. The presence of a TRAF-like domain in the XAF1 N-terminal domain further supports that XAF1 could play a role in NF-κB-mediated immunity. In fact, it has been reported that XAF1 regulates TNF signaling by inhibiting NF-κB activation (9). Moreover, it was reported that better control viremia rebound after antiretroviral therapy interruption in HIV patients was associated with upregulation of XAF1 transcripts (10). Additionally, whole transcriptome analysis from both COVID19 and influenza patients unveiled XAF1 as a common upregulated gene signature (11). Furthermore, recent publications pointed to XAF1 as playing an important role against virus infections (12, 13). Even though from cancer field XAF1 is considered a master regulator in cell death, the impact of IFN-dependent XAF1 induction upon virus infection has been poorly studied.

Using a Sterile alpha and TIR motif containing 1 (SARM1) knockout mouse model, we previously suggested *Xaf1* as a candidate gene to be involved in viral phenotypes previously attributed to SARM1 deficiency, including infections in the central nervous system (CNS) (14). SARM1-deficient mice generated in a 129 genetic background and backcrossed in C57BL/6J mice (SARM1^*AD*^) (15) harbor a XAF1 polymorphism not present in wild type (WT) C57BL6/J mice (from 129 origin) due to the chromosomal proximity of *Sarm1* and *Xaf1* genes. These SARM1-deficient mice showed significant reduction of *Ccl3, Ccl4* and *Ccl5* expression (15). However, when C57BL/6J CRISPR-Cas9 SARM1-KO mice (SARM1^*AGS*^) were generated, no significant decrease of these cytokines was observed (14). Additionally, the protective phenotype attributed to SARM1-deficiency upon VSV infection observed in SARM1^*AD*^ mice, was not observed in SARM1^*AGS*^ mice. Taken together, these results suggested the possibility that *Xaf1*, with different polymorphisms in C57BL/6J and 129 mice, may be responsible for the phenotypes observed in SARM1^*AD*^ but not in SARM1^*AGS*^. In an attempt to study the relevance of XAF1 in antiviral immunity and discern the viral phenotypes associated with SARM1-deficiency, we generated a CRISPR-Cas9 XAF1 C57BL/6J knockout (XAF1-KO) mouse and found that XAF1-KO mice have increased susceptibility to VSV infection, supporting the protective phenotype observed in SARM1^*AD*^ mice, whilst having limited impact on viral replication. In addition, XAF1-KO mice showed increased cytokine and proinflammatory responses upon VSV infection as well as augmented CNS inflammation, immune cell infiltration and disease exacerbation. Furthermore, RNAseq data unveiled both a dysregulated immune response and diminished capacity to restore homeostasis after infection in XAF1-KO mice. Overall, these results suggested that the phenotypes observed in SARM1^*AD*^ and not in SARM1^*AGS*^ mice were not likely attributed to XAF1 polymorphism and suggest XAF1 to be a protective factor against viral-induced neuropathogenesis.

## Results

### XAF1 deficient mice showed increased susceptibility to severe infection with VSV and WNV neurotropic viruses and normal responses to IAV respiratory infection

Previously, phenotypic characterization of SARM1-deficient C57BL/6J mice harboring XAF1 from 129 origin suggested a role of XAF1 in CNS viral infections (14) To address such a possibility, we generated a XAF1-KO mouse using CRISPR-Cas9 technology. Since commercially available antibodies failed to detect XAF1 by western blot, validation of the model was carried out by RT-qPCR in which *Xaf1* transcripts were detected 24 h after Poly (I:C) administration of WT mice but significantly diminished in XAF1-KO mice (Supplemental Fig. 1). To test the effect of XAF1 deficiency on neurotropic viral infection, groups of mice were infected with different infectious doses of VSV (high dose: 10^7^ PFU; middle dose: 10^5^ PFU and low dose: 10^3^ PFU). Upon VSV infection, XAF1-KO mice showed increased mortality rates in both the 10^7^ and 10^5^ PFU groups reaching 100% lethality by day 9, while XAF1-KO mice infected with the low dose reached 40% mortality (Fig. 1A). Strikingly, the middle dose was enough to cause 100% lethality in the XAF1-KO while in WT mice, mortality was significantly reduced to 50%. To evaluate the pathological implications of the infection in absence of XAF1, mice were infected with 10^5^ PFU of VSV and brain samples were harvested at days 2 and 5 post infection. Viral titers were determined by plaque assay and viral replication was assessed by RT-qPCR. Interestingly, no significant differences were observed between WT and XAF1-KO mice in terms of infectious VSV particles or RNA copies at both days tested (Fig. 1B and Supplemental Fig. 2). Similarly, with the exception of a slight but significant increase at day 5 postinfection in XAF1-KO mice, viral growth in the olfactory bulb (OB), which is primary site of VSV replication *in vivo*, was similar between WT and XAF1-KO animals (Supplemental Fig. 2). Additionally, footpad infection of XAF1-KO mice with another neurotropic virus (WNV) showed a slight increase in disease severity compared to that of WT mice, although it did not reach statistical significance (Fig. 1C). Interestingly, when a respiratory virus was used, i.e. IAV, there was no difference in morbidity or mortality between WT and XAF1-KO mice. Additionally, no difference in viral lung titers for IAV were observed either at day 2 or day 6 pi (Supplemental Fig. 3). These results suggest XAF1 as a protective factor against tissue inflammation in neurotropic virus infection with limited antiviral role against IAV or WNV *in vivo*.

**Figure 1:**
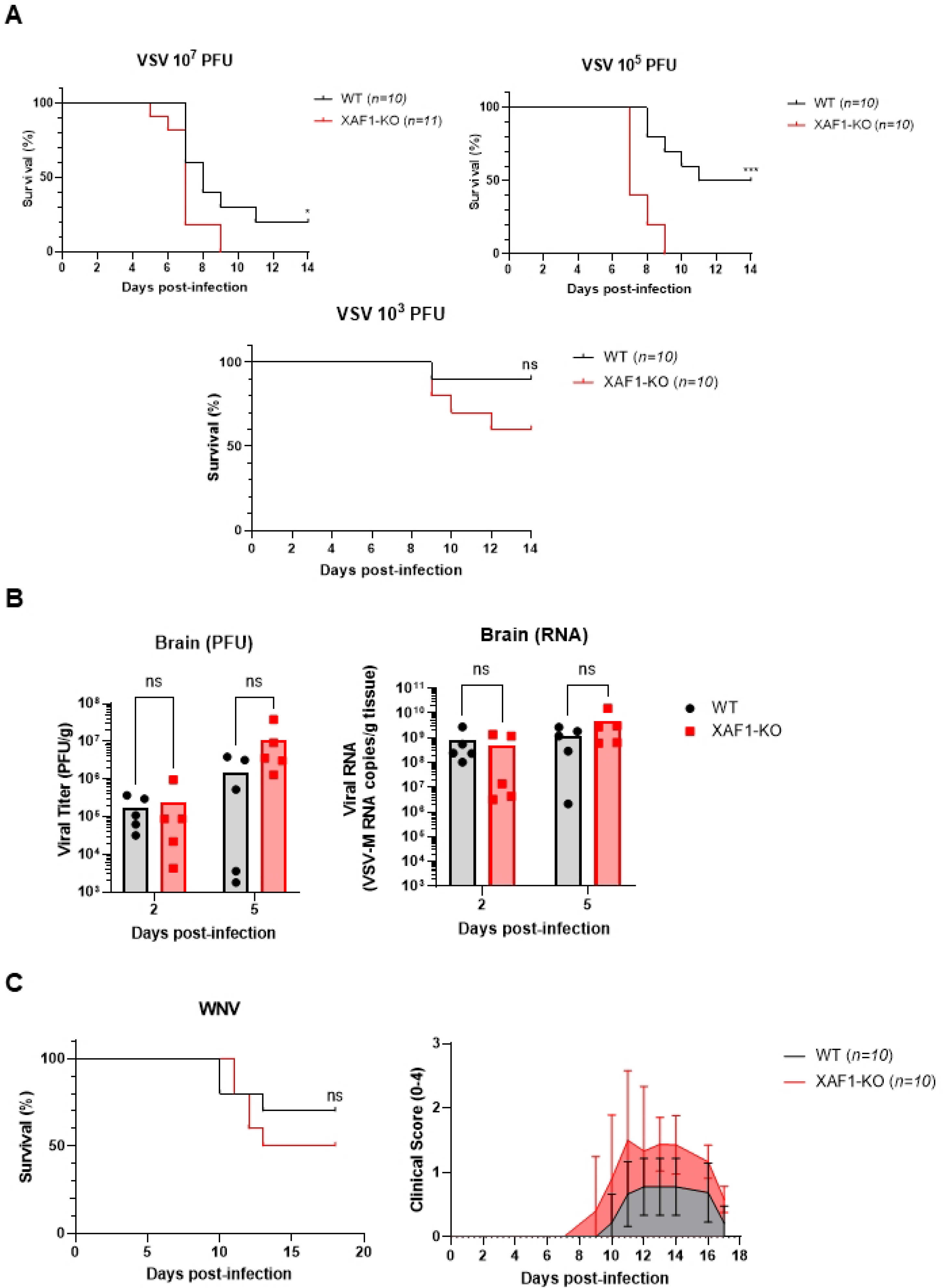
XAF1-KO mice are more susceptible to fatal VSV infection. A) Groups of mice were anesthetized with ketamine/xylazine as described in materials and methods and intranasally infected with the indicated doses of VSV and mortality were observed for 14 days. B) Brain samples were collected at days 2 and 5 post-infection and viral titer was determined by plaque assay and viral RNA was determined by VSV-specific qPCR (see primers on Table S1). C) Groups of mice were anesthetized as above and infected in the footpad with 100 PFU of NY/99 strain of WNV. Mortality was observed for 18 days and clinical signs were quantified as described in (41) (0: healthy; 1: ruffled hair, lethargy, hunched posture, no paresis, normal gait; 2: altered gait, limited movement in 1 hind limb; 3: lack of movement, paralysis in 1 or 2 hind limbs; 4: moribund).Data is representative of 3 independent experiments and each dot represents one individual animal. Data is presented as bars. ns denotes no statistically significant differences.

### XAF1 deficiency leads to an increase in markers of inflammation upon VSV infection in the brain

To further understand the cause of increased morbidity and mortality in XAF1-KO mice when infected with VSV, we next examined levels of inflammatory markers in the CNS during infection. To that end, WT or XAF1-KO mice were mock infected or infected with VSV at 10^5^ PFU. Brain samples were harvested at day 5 post-infection and expression of surrogate markers for tissue inflammation were analyzed by RT-qPCR. Compared to mock-infected controls, both WT and XAF1-KO mice showed augmented levels of pro-inflammatory cytokines and chemokines, including *Ifnβ, Il1β, Il6* and *Ccl2* (Fig. 2A), following a trend amplified in absence of XAF1. Additionally, flow cytometry analysis was performed to identify immune cell populations in the brain of infected animals at 5 dpi. As shown in Fig. 2B and 2C, an increase in CD45^high^ CD11b^high^ cells was observed in XAF1-KO mice. Of note, XAF1-KO mice showed a trend towards reduction of the resident activated microglia population, defined as CD45^low^ CD11b^high^ subset, upon infection. No major changes were observed in the resting microglia subset (CD45^low^ CD11b^low^) (Fig. 2C). Overall, the data indicated that XAF1 deficiency led to an abnormal immune response in the brain upon viral infection, characterized by increased inflammatory markers and immune cell infiltration.

**Figure 2:**
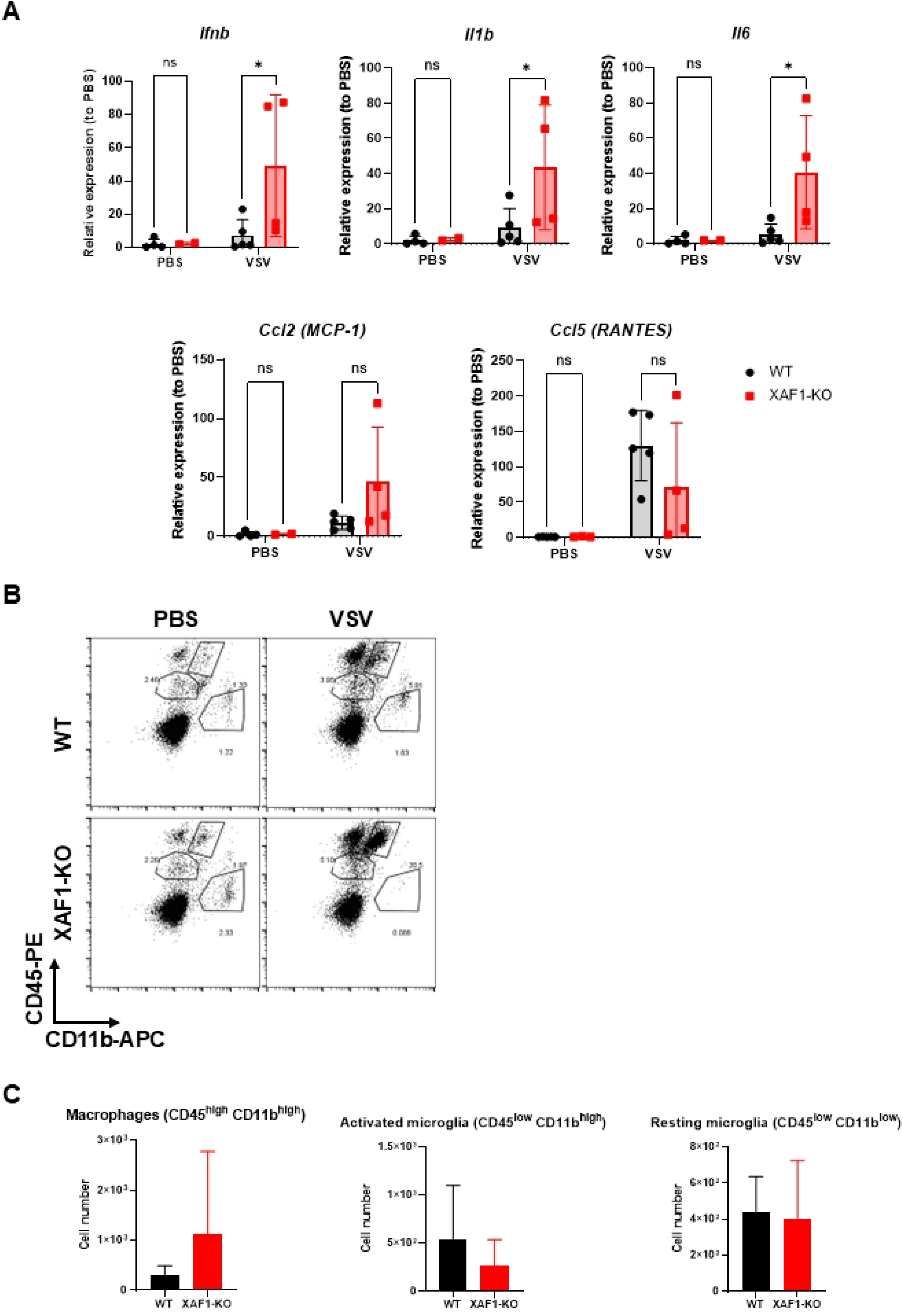
Aberrant immune response upon VSV infection in XAF1-KO mice. A) Groups of mice were mock-infected or infected with VSV (10^5^ PFU) and total RNA was extracted from brain samples on day 5 post infection. Levels of proinflammatory cytokines were measured by qPCR (see primers on Table S1). B) Flow cytometry analysis of brain samples from mock-infected or VSV-infected mice on day 5 post-infection. Groups of mice (*n=3*) were sacrificed, perfused with PBS and individual cells were obtained for antibody staining for FACS analysis. CD45-PE, and CD11b-APC antibodies were used. Quantification of total positive stained cells is shown in (C). Each dot denotes an individual animal and data is presented as mean ± SD.

### CNS inflammation and viral-induced neuropathogenesis is exacerbated in XAF1-deficient mice

To further evaluate the role of XAF1 on CNS inflammation and pathogenesis exerted by VSV infection, brain samples of mice mock infected or infected with VSV, were harvested at 6 dpi, formalin-fixed, processed and subjected to hematoxylin & eosin (H&E) staining. While WT infected mice showed moderate inflammation and immune cell infiltration, this was exacerbated in XAF1-KO mice, which also showed increased perivascular cuffing in the OB (Fig. 3A). In addition, viral-induced encephalitis, vessel thickening, and necrosis were extensively observed in XAF1-KO mice. A quantification of pathological observations revealed that XAF1-KO mice presented with higher clinical scores and worse disease in comparison to WT mice (Fig. 3B). Furthermore, immunohistology analysis unveiled extensive viral antigen in neurons in the OB at day 6 post infection, overlapping with areas of viral-induced neuropathogenesis and tissue damage of the CNS (Fig. 3C). These findings support the protective role of XAF1 during viral-induced tissue damage and CNS disease by preventing exacerbated viral-induced encephalitis.

**Figure 3:**
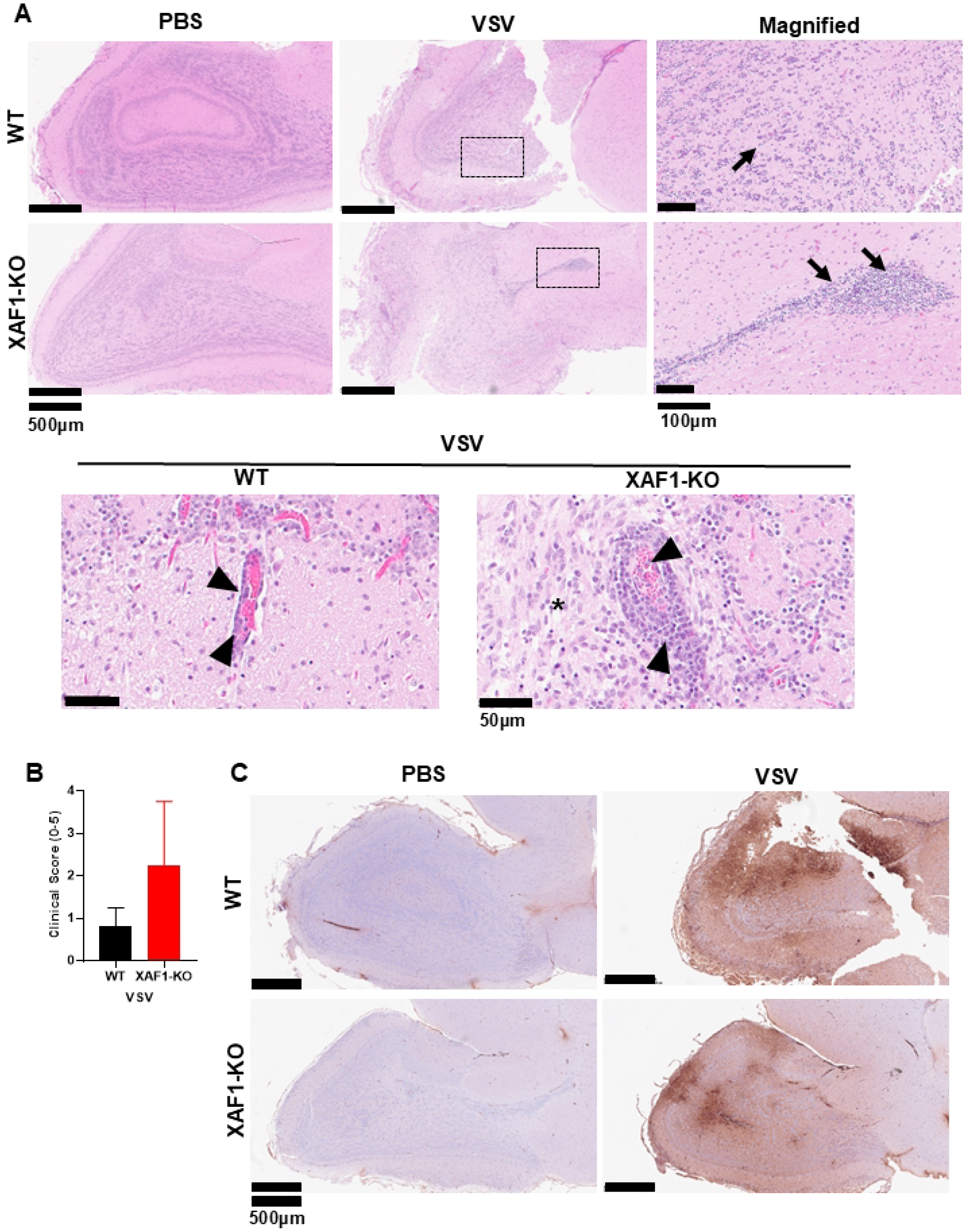
Viral-induced neuropathogenesis is increased in XAF1-KO mice. A) Hematoxylin & eosin staining of brain samples on day 5 post infection. Olfactory bulb sections are shown. Cell infiltration and encephalitis are shown by arrows, arrow heads indicate thickening of vascular endothelium and asterisk edema (scale bar is indicated as horizontal bar). Magnified sections correspond to areas marked with the black rectangle. B) Clinical score (0-5) from each group of infected mice. Score was assigned as follows: 0: no lesions; 1: minimal perivascular inflammatory infiltrate, no gliosis, no neurodegeneration, no satellitosis and no necrosis; 2: mild perivascular inflammatory infiltrate, mild gliosis, no neurodegeneration, no satellitosis and no necrosis; 3: Mild perivascular infiltrate, choroiditis, mild gliosis, neurodegeneration, satellitosis and necrosis; 4: Moderate perivascular and brain parenchyma inflammatory infiltrate, choroiditis with moderate gliosis, mild neurodegeneration, mild satellitosis and mild necrosis; 5: Severe parenchymal inflammatory infiltrate with moderate to severe gliosis, neurodegeneration, satellitosis and necrosis. C) Immunohistological VSV-M staining of olfactory bulb from mice in A). Scale bars are indicated in each case with a horizontal bar.

### Transcriptomics analysis of the CNS unveils both dysregulated proinflammatory profile and tissue remodeling impairment in XAF1-deficient mice upon VSV infection

To have a better understanding of both the breadth and magnitude of the host response in the CNS to VSV, bulk RNAseq analysis was performed and the differential gene expression profile was analyzed in WT and XAF1-KO infected mice. Transcriptome analysis revealed a total of 462 and 215 genes that were significantly upregulated and downregulated respectively (Fig. 4A). Compared to WT-infected mice, an upregulation of genes involved in response to inflammation and tissue injury, such as *Saa1* as well as pro-inflammatory cytokines-related genes such as *Ccl7* and *Il6 as well as* IFN response (*Ifna6, Ifnab*) were found in XAF1-KO mice, confirming the exacerbated immune response in these mice. On the other hand, genes involved in tissue remodeling and cell death such as *Ms4a15, Col1a1* and *Bglap* were significantly downregulated (Fig. 4A). Moreover, gene ontology analyses revealed an enrichment of a variety of signaling pathways, particularly related to innate immunity, cellular response to Ifnb, Il1b signalling pathway and inflammatory responses in XAF1KO-infected mice compared to WT-infected mice (Fig. 4B). Overall, transcriptomic data suggested both an exacerbated innate immune response and a dysregulated capacity to recover homeostasis after viral infection of the CNS.

**Figure 4:**
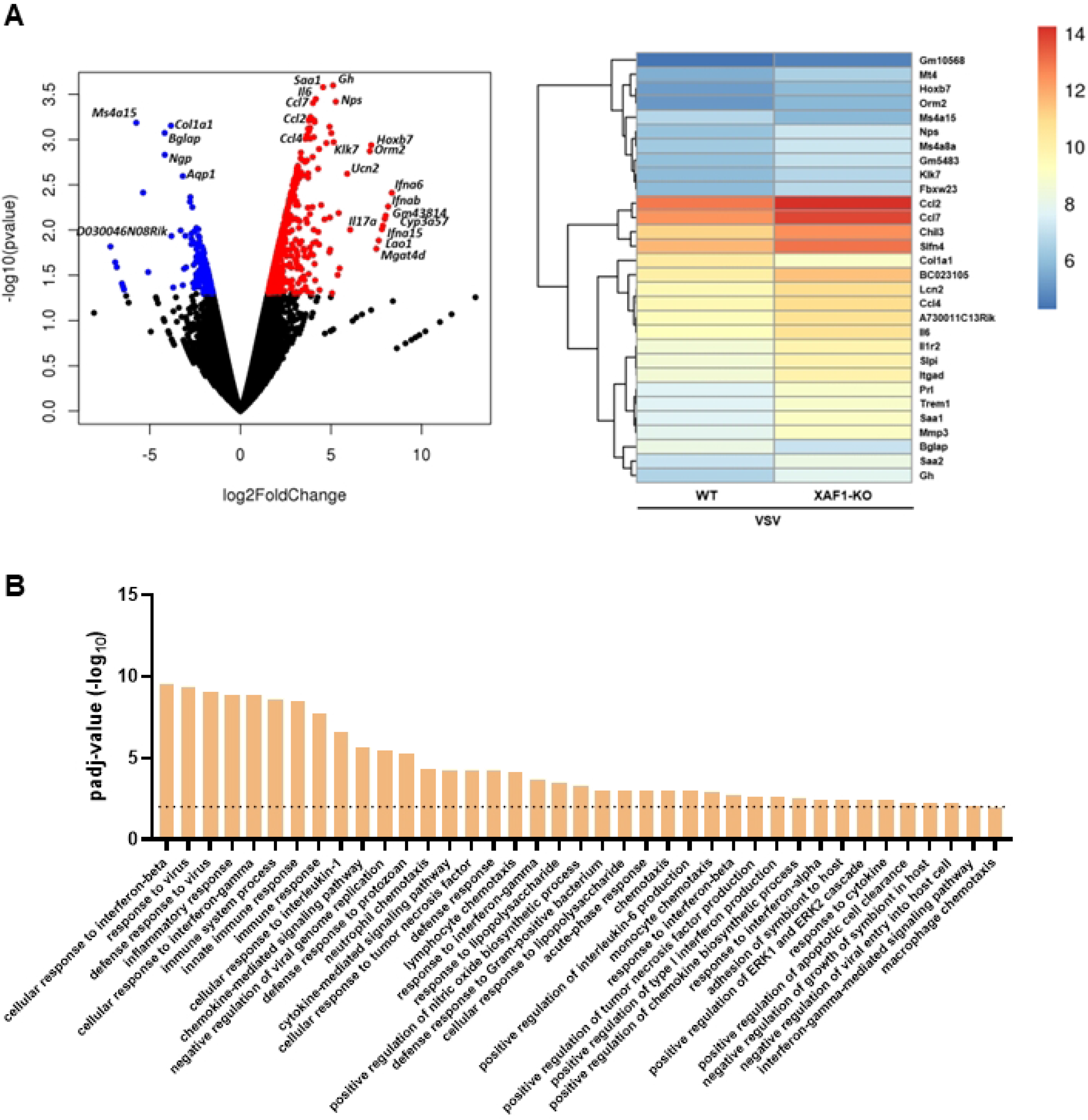
XAF1 deficiency unveils an unbalanced host response against neurotropic virus infection. A) Volcano plot and heatmap plot showing differential gene expression analysis of WT vs XAF1KO VSV-infected mice. Upregulated and downregulated genes are shown in red and blue respectively. B) Gene ontology data showing statistically significant increased (*p*≤0.01) cellular pathways and biological processed in XAF1-KO infected mice compared to WT-infected mice. The dotted line indicates *p*≤0.01.

### XAF1 deficiency leads to an increased IFN response

To better understand the implications of XAF1-deficiency in disease severity observed in mice, we studied the IFN response in XAF1-deficient cells *in vitro*. To this end, WT or XAF1-KO 3T3 cells were treated with 1000 U of universal IFN for 24h and *Mx1* expression was measured by qPCR as an indicator of ISG induction. XAF1-deficient 3T3 cells showed a strong increase in *Mx1* expression upon IFN stimulation in comparison to WT controls (Fig. 5A). To further elaborate on elevated ISG expression, cells were incubated with different PAMPs, i.e. Poly IC, HT-DNA and LPS for 8h and transcript levels of *Mx1* and *ISG15* were measured. Similarly, XAF1-KO cells in comparison to WT equivalents, had a significant increase in both *Mx1* and *ISG15* expression upon PAMPs-stimulation (Fig. 5B). These results suggest a dysregulated IFN response in XAF1-deficient cells upon external stimuli, indicating a negative regulatory role for XAF1 in IFN response. To confirm this hypothesis, 293T cells were transfected with increasing amounts of plasmid encoding for XAF1 together with a plasmid containing a Firefly luciferase cassette under the control of IFN-sensitive response element (ISRE) promoter and a plasmid encoding for Renilla-luciferase as constitutive control. Upon IFN treatment, luciferase activity was decreased in parallel to increasing XAF1 concentrations (Fig. 5C). In conclusion, results suggest a role of XAF1 in modulating IFN signaling *in vitro* and XAF1 as a negative regulator of the IFN response.

**Figure 5:**
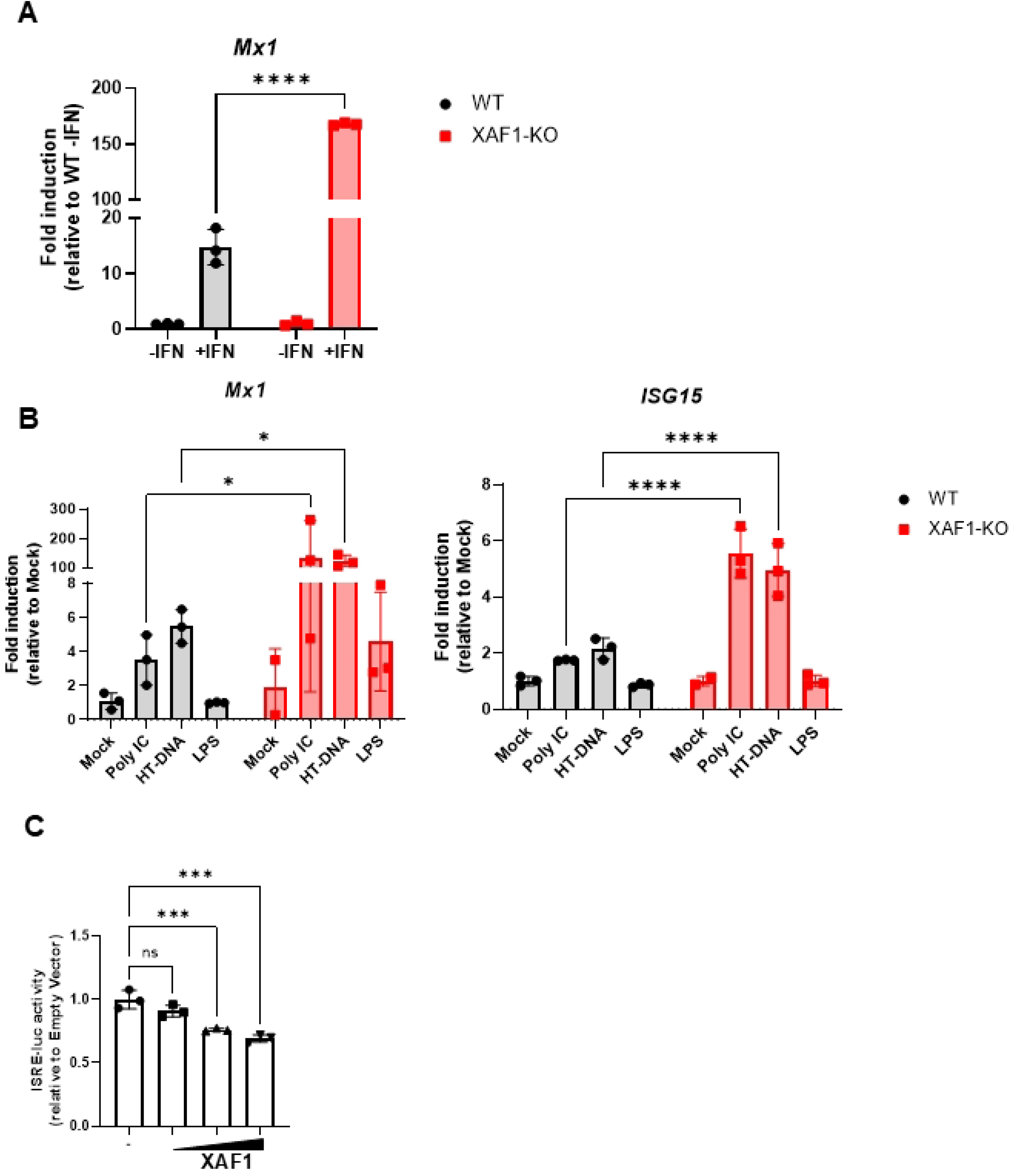
XAF1 deficiency increased IFN response *in vitro*. A) 3T3-WT or XAF1-KO cells were mock-treated or treated with 1000 U of universal IFN for 24h and *Mx1* expression was measured by qPCR. B) 3T3 cells were treated with Poly (I:C) (1 µg/ml), HT-DNA (10 µg/ml) or LPS (10 µg/ml) for 8h and *Mx1* and *ISG15* were measured by qPCR using the primers listed in Table S1. Data is presented as fold induction over unstimulated cells. C) 293T cells were co-transfected with ISG54 plasmid (50 ng), pTK renilla luciferase (20 ng) and pCAGGs-XAF1 (2 ng, 20 ng and 200 ng). At 24h, cells were treated or not with 1000 U IFN and luciferase activity was measured at 24h. Data is presented as an average of three independent biological replicates ± SD. Statistically significant differences were obtained after 2-way ANOVA and denoted as asterisks (\*\*\*\**p*≤0.0001, \*\*\**p*≤0.001, \**p*≤0.05). Non-IFN-treated cells signal was subtracted as background.

## Discussion

Mostly known from cancer field, XAF1 has been suggested as a candidate gene in virus infections due to its IFN responsiveness and its role in IFN-induced apoptosis (16, 17). CRISPR-Cas9 generated C57BL/6J SARM1-KO mice failed to recapitulate the viral phenotypes and chemokine production observed in mice lacking SARM1 and containing neighboring genes from 129 origin, including *Xaf1, Ccl3, Ccl4* and *Ccl5* (MIP locus), (14). Among all these genes, the only one that showed substantial isoform polymorphism was *Xaf1*. This prompted us to examine the role of this factor in chemokine expression and protection from severe VSV neuropathogenesis. In fact, recent studies further supported a protective role of XAF1 against infection with a variety of RNA viruses (12, 13). *Xaf1* polymorphism between 129 and C57BL/6J backgrounds included several SNPs, a deletion of a 248 bp in exon 6 in 129 mice and differences in alternative splicing of the different transcripts giving rise to novel isoforms previously uncharacterized (14). Thus, differences in phenotypes found in Sarm1^*AD*^ mice from 129 origin versus C57BL/6J SARM1-KO could be due to genomic differences in *Xaf1*. Nevertheless, one could not exclude that differences in expression of chemokines, including *Ccl3, Ccl4* and *Ccl5* genes, also present in the same 129 mice-derived locus present in Sarm1^*AD*^ mice, were playing a role.

The results obtained here support the hypothesis of a protective role of XAF1 in CNS viral infections. Interestingly, protection was not due to a restriction on viral replication. Rather, it was mediated by reduced expression of proinflammatory cytokines, since XAF1-KO mice tended to show increased levels of *Ifnb, Il1b, Il6* and *Ccl2*. These results agree with previous studies suggesting that protection in the CNS is mediated by a controlled immune response, rather than viral control (15, 18). Interestingly, the impact of XAF1 on gene expression was specific for a subset of cytokines, as we previously have shown that neither *Ccl3, Ccl4* nor *Ccl5* expression was modulated by XAF1 (14). Protection against other neurotropic viruses like WNV was also observed although to a lesser extent. Mediators controlling different viruses such as key chemokines involved in WNV infection could explain the differences observed. Ccr5, a chemokine receptor that interacts with different ligands, including Ccl3, Ccl4 and Ccl5, is a key chemokine in controlling WNV infection (19). In fact, results presented here showed that *Ccl5* was not upregulated in XAF1-KO mice upon VSV infection, suggesting that chemokines controlling WNV and VSV may differ, and Ccl5 could be a possible key candidate chemokine for WNV control, explaining the differences between these two viruses in XAF1-mediated susceptibility to severe infection. Also, differences between infection model *ie*, intranasal for VSV *vs* footpad injection for WNV should be considered when comparing differences in susceptibility.

Here, we also show that XAF1 plays a limited role in respiratory virus infection and lung pathology. These results contrast with those found by Han et al (12), which suggested that XAF1 not only prevented lung inflammation but could also directly impact IAV infection. In our studies, no differences were found in IAV replication between wild type and XAF1-KO mice. Differences in the genetic background, the way the knockout was designed and sequence targeting could be among the reasons for this discrepancy.

Immune cell infiltration into the CNS after infection was increased in XAF1-KO mice. Infiltrated macrophages are key cells involved in inflammation and secretion of proinflammatory cytokines that can exacerbate pathology and disease, including infections in the CNS (20). Here, we observed that macrophages in the CNS were more abundant in XAF1-KO mice upon VSV infection. Ccl2, a key ligand involved in monocyte release from the bone marrow into the blood (21, 22), was increased in XAF1-KO deficient mice, correlating with the augmented levels of macrophages invading the CNS. Many studies have previously shown increased immune cell infiltration upon acute viral infection (23-25). High inflammatory macrophages (Ly6C^high^) are thought to be the main Ccl2 producers (26, 27) and even though it has been described that both neurons and microglial cells promote cytokine production upon viral infection (15, 28), further studies will be required to better understand the source of these cytokines. Microglia cells are specialized immune cells residing in the CNS that protect against a variety of viral infections, including WNV (29), JEV (30) and VSV (31, 32). The results shown here indicate a trend towards decrease of microglia activation in XAF1-KO mice during acute viral infection. It has been suggested that activation of microglia occurs upon neural damage and is regulated by IFN produced in neurons and astrocytes (31). In fact, protection against virus invasion into the CNS has been related to microglia activation and T-cell activation (32, 33). This goes in line with our results shown here, where activation of microglial cells was hampered in XAF1-KO mice, hence failing to protect against severe pathology. Whether there is a link on XAF1-mediated microglia activation is something that would require further study. Gene ontology analysis unveiled that XAF1-KO mice showed a significant dysregulation of signaling pathways including cellular response to Il1b. It is known that Il1b production promotes the disruption of blood-brain barrier (BBB) integrity and increases neuroinflammation (34, 35). With the results provided here, it is possible that the increased pathology observed in XAF1-KO mice upon VSV infection is related to the alteration and increased permeability of the BBB. This BBB dysfunction has been described upon infection with different neurotropic viruses, facilitating both immune cells and cytokine infiltration into the CNS (36). However, the relevance of XAF1 and BBB integrity upon virus infection in the CNS remains unclear. Although we observed increased IFN upon infection in XAF1-KO mice, only a subset of ISGs were upregulated, indicative of an unbalanced IFN response in XAF1-KO mice that might lead to a failed activation of microglial cells. This phenotype was also found in XAF1-deficientcells, where *Mx1* and *ISG15* transcripts were upregulated upon IFN treatment and different immune sensor agonists, suggesting a role of XAF1 in regulating the response to IFN. It has been suggested that XAF1 is required for a IRF1-dependent innate immune response (1, 12). Our results go in line with those found by other authors describing that both, *Mx1* and *ISG15* are IRF1-targeting ISGs (37, 38), also since stimulus-dependent gene expression differences were observed, future studies will be needed to discern the molecular mechanisms underlying the interplay between XAF1 and IFN response. In any case, our results indicate that XAF1 expression protects from viral-induced neuropathology, and points to this gene as a regulator of exacerbated immune-driven inflammation in the CNS.

## Materials and Methods

### Cell lines

HEK293T (ATCC, CRL-3216) and 3T3 cells (ATCC, CRL-1658) were maintained in Dulbecco’s modified Eagle’s medium (DMEM) (Corning) supplemented with 100 U/ml of streptomycin/penicillin (Corning), 2 mM L-glutamine (Corning) and 10% fetal bovine serum at 37ºC and 6% CO2. 3T3 XAF1-KO cells were already described in (14).

### Mice

XAF1-KO mice on C57BL/6J background were generated using the CRISPR design tool (genome-engineering.org) to select the guide sequences AGCTTCCTGCAGTGCTTCTGTGG and AGGCTGACTTCCAAGTGTGCAGG that were cloned into pSpCas9n(BB)-2A-GFP as described (39). The resulting plasmid was injected into male pronuclei of C57BL6/J mouse embryo as described in (14). Mice were genotyped by PCR using the primers listed in Table S1 and housed in a pathogen-free barrier facility and maintained with food and water *ad libitum*.

### Viral infections in mice

6-8 week old mice were anesthetized with ketamine/xylazine (95 mg/kg ketamine and 10 mg/kg xylazine) and infected intranasally with 30 µl PBS containing the indicating doses of VSV or 100 PFU of Influenza A/PR/8/34 (H1N1) (PR8) virus. For WNV infection, 100 PFU of WNV NY/99 was administered by infection in the footpad. Animals were humanely sacrificed when reached the humane endpoint (loss of 25% initial body weight or severe paralysis). For tissue harvesting, animals were injected intraperitoneally with sodium pentobarbital (150 mg/kg) and tissue samples were harvested at the indicated time points. Virus titers recovered from infected mice were determined by plaque assay. Animal studies were approved by the Institutional Animal Care and Use Committee of Icahn School of Medicine at Mount Sinai.

### Lung pathology and immunohistochemistry

Tissue samples of infected mice were harvested at the indicated time points and fixed in a PBS solution containing 10% formalin. Parafin-embeded brain sections were either stained with hematoxylin and eosin or incubated with anti-VSV M antibody (Kerafast) followed by pathology scoring by the Neuropathology Brain Bank and Research Core and Department of Pathology at Icahn School of Medicine at Mount Sinai.

### RT-qPCR

Total RNA was extracted using the kit EZNA total RNA kit and RNase-free DNase (Omega). RNA was first reversed transcribed using Maxima Reverse Transcriptase and oligo-dT (ThermoFisher). qPCR was performed using the resulting cDNA and the primers listed in Table S1. Data is shown as relative expression relative to mock.

### IFN treatment and PAMP stimulation

3T3 WT and XAF1-KO cells were washed with PBS and incubated with DMEM containing or not 1000 IU of universal IFN (consisting of recombinant human interferon alpha A and alpha D) (uIFN) (pbl bioscience). Additionally, DMEM containing Poly(I:C) (InvivoGen) (1 µg/ml), HT-DNA (Sigma) (10 µg/ml) or LPS (Enzo) (10 µg/ml) for 8h. Then, cells were washed, and total RNA was extracted as described above.

### Flow cytometry

VSV-infected mice were sacrificed at day 5 pi and perfused three times with 30 ml of sterile PBS. Single cell suspensions were prepared as described (40). Briefly, brain samples were harvested, cut and incubated in 2 ml of digest solution (RPMI 10% FBS 0.2 mg/ml collagenase IV (Sigma), 0.05% DNase I (Roche)) for 1h at 37°C. Then, digested tissue was dissociated with a syringe and passed through a 70 µm cell strainer into a 50 ml tube and washed three times with PBS-EDTA 0.05%. The pellet was resuspended in 40% Percoll and layered on a 15 ml tube containing 80% Percoll. Cells were washed three times with FACS buffer (0.5 mM EDTA, 0.5% BSA) and stained using eViability e450 (BD), CD45-PE (BD) and CD11b-APC (BD) antibodies. After washing three times, cells were resuspended in 300 µl of FACS buffer and passed through a Gallios Cytometer (Beckman) and data were analyzed using the software FloJov_10.6.2.

### Luciferase assay

For luciferase assays, HEK293T cells were transiently transfected with pRL-TK and ISG54-luciferase vectors along with the plasmid pCAGGs-XAF1. At 24 h after transient transfection, cells were treated overnight with universal IFN type I (1,000 IU/mL) (PBL) and luciferase activity was measured using the Dual-Luciferase Assay System (Promega) according to the manufacturer’s instructions. Firefly luciferase values were normalized to Renilla, and the fold induction was calculated as the ratio of IFN-stimulated versus unstimulated cells.

### RNAseq

RNA from brain samples from VSV-infected mice at day 5 pi were extracted as above. Poly-A selection-based mRNA enrichment, random priming with subsequent cDNA synthesis and Illumina technology-based sequencing of 2×150bp read length were performed by GENEWIZ (South Plainfield Overed, NJ).

## Acknowledgments

We thank all past and current members of the A.G.S. laboratory for their technical and theoretical support, in particular Richard Cadagan, Ashely Diaz-Tapia and Daniel Flores. We also thank members of the AGS laboratory for their insights into the results provided in this manuscript. We acknowledge Claudia Sanctis and Diana Dangoor at the Neuropathology Brain Bank & Research Core and Ma Gonzalez, Ying Dai and Monica Garcia-Barros from the Comparative Pathology Laboratory at ISSMS.

## Figure legends

**Figure S1: XAF1-KO mice validation.** WT and XAF1-KO mice were intravenously administrated with 100 µg of Poly (I:C) and total RNA from spleen samples were harvested at 24 h and XAF1 transcripts were analyzed by qPCR using the primers listed on Table S1.

**Figure S2: XAF1 deficiency does not affect VSV replication *in vivo*.** Mice were intranasally infected with 10^5^ PFU of VSV and olfactory bulb (OB), brain and lung samples were harvested at days 2 (A) and 5 pi (B). Data is presented as mean ± SD. 2-way ANOVA was used with Sidak correcting test for multiple comparisons. Statistical differences are denoted with asterisks (** p<0.01).

**Figure S3: XAF1-KO mice do not display IAV phenotype.** A) Groups of WT (*n=9*) and XAF1-KO mice (*n=11*) were infected with 100 PFU of IAV (A/Puerto Rico/8/1934 (H1N1)) and weight loss and survival were monitored for 14 days. B) Lung samples were harvested at days 2 and 6 pi and viral titer was determined by plaque assay. Data is presented as mean ± SD. 2-way ANOVA was used with Sidak correcting test for multiple comparisons (ns = no significant).

